# Cellular Composition Accounts for Much of the Bulk Glutamine–IFN-γ Transcriptional Axis in Melanoma: A Cross-Cohort and Single-Cell Analysis

**DOI:** 10.1101/2025.09.28.679008

**Authors:** Islam Asal

## Abstract

**Background:** An inverse relationship between glutamine-associated and interferon-γ (IFN-γ)-associated transcription in bulk melanoma could indicate metabolic antagonism or simply different cellular origins. We tested these alternatives and evaluated prognostic and immunotherapy-response associations.

**Methods:** Continuous GSVA scores were analyzed in TCGA-SKCM (469 tumors), an independent melanoma survival cohort (GSE65904; 214 tumors), a pretreatment nivolumab cohort (GSE91061; 49 tumors), and single-cell RNA sequencing from 19 patients (GSE72056; 4,645 cells). Cox models used harmonized endpoints and clinical covariates. Bulk composition was examined with ABSOLUTE purity, EPIC, and MCP-counter. Single-cell scores were summarized by patient and compartment; paired tests treated patients, not cells, as independent units. Response models used Firth logistic regression, leave-one-out discrimination, and simulation-based precision analysis.

**Results:** In TCGA, the candidate scores were inversely correlated before adjustment (r=−0.397; 95% CI −0.470 to −0.318) but progressively attenuated after purity/sample-type adjustment (r=−0.218), EPIC adjustment (r=−0.146), and MCP-counter adjustment (r=0.019). Single-cell analysis localized higher glutamine-associated scores to malignant cells than paired T, NK, or B-cell compartments, while IFN-γ-associated scores were lower in malignant cells than paired T-cell and macrophage compartments. IFN-γ was associated with lower mortality in TCGA (adjusted hazard ratio [HR] per SD 0.648; 95% CI 0.555–0.757) and GSE65904 (HR 0.667; 95% CI 0.534–0.835). Glutamine-associated scores were not independently prognostic in TCGA (HR 0.966; p=0.673) and were uncertain in GSE65904 (HR 0.823; p=0.084). The interaction was null in TCGA, nominal for the candidate sets in GSE65904, and null with curated sets. No response association was supported in GSE91061; approximately an odds ratio of 3.25 per SD was required for 80% simulated power.

**Conclusions:** Cellular composition accounts for much of the bulk glutamine–IFN-γ transcriptional axis in melanoma. IFN-γ-associated transcription is a reproducible prognostic correlate, but glutamine-associated transcription provides no stable incremental outcome information. These data do not establish metabolic competition or treatment prediction; spatial or functional measurements and larger treatment-comparative cohorts are needed.

## Introduction

Melanoma cells can depend on glutamine-derived carbon and nitrogen, while activated lymphocytes require coordinated nutrient availability to sustain effector function. Experimental studies therefore make metabolic competition within the tumor microenvironment biologically plausible [1–3]. IFN-γ-responsive transcription marks antigen presentation, chemokine signaling, cytotoxic inflammation, and adaptive immune resistance, and related expression profiles have been associated with benefit from PD-1 blockade [4].

Bulk tumor transcriptomes, however, combine malignant, immune, stromal, and vascular signals. Immune-enriched transcription is associated with melanoma outcome [5], and single-cell studies show marked interpatient and intratumoral heterogeneity [9]. A negative bulk correlation between glutamine-associated and IFN-γ-associated scores could therefore arise from a biological interaction within cells, from variable proportions of cell populations that express the programs differently, or from both. Bulk correlation alone cannot distinguish these explanations.

Our preliminary analysis proposed a median-defined four-group classification. Subsequent methodological review identified limitations in endpoint harmonization, small-sample response analysis, incremental-value assessment, and treatment of composition. More importantly, the original bulk-only analysis could not localize the two programs to cellular compartments.

We therefore performed a strengthened, cross-cohort analysis with four complementary datasets. We asked whether (i) the bulk inverse association persisted after several composition adjustments, (ii) malignant and immune compartments showed different score localization at single-cell resolution, (iii) prognostic associations replicated in an independent survival cohort and with curated signatures, and (iv) pretreatment scores showed sufficiently precise evidence of immunotherapy-response association. The primary estimands used continuous scores; median groups were descriptive only.

## Methods

### Study design and data sources

This retrospective secondary analysis used public, deidentified transcriptomic and clinical data. Four cohorts served distinct roles: TCGA-SKCM for discovery survival and deconvolution; GSE65904 for independent survival validation; GSE91061 for exploratory pretreatment response analysis; and GSE72056 for single-cell localization. Because the treated dataset lacks a non-immunotherapy comparator, response associations are prognostic-within-treated-cohort associations and cannot establish treatment-predictive utility.

### Bulk melanoma cohorts

TCGA-SKCM RNA-sequencing and clinical data were processed as described in the corrected reanalysis [5–7]. One expression profile per patient was retained, giving 469 patients. Overall survival (OS) time, event status, age, sex, and AJCC stage were obtained from the TCGA Clinical Data Resource [7]. The primary multivariable set contained 417 patients and 194 deaths. Stage was grouped as 0/I/II versus III/IV. ABSOLUTE purity and sample type were included in sensitivity analyses [12].

GSE65904 is an independent Illumina HumanHT-12 v4 melanoma expression cohort [8]. Probe-level expression was mapped through GPL10558; multiple probes mapping to one gene were averaged. Disease-specific survival (DSS), age, sex, stage, and tissue source were harmonized before modeling. Of 214 available tumors, 210 had eligible DSS data and 203 had complete model covariates, including 99 disease-specific deaths.

GSE91061 contains pretreatment melanoma biopsies from patients receiving nivolumab [6]. Complete or partial response was classified as response; stable or progressive disease as non-response; unknown response was excluded. Forty-nine pretreatment samples remained, including 10 responders. The analysis does not compare nivolumab with another treatment.

### Single-cell cohort and annotation

GSE72056 contains single-cell RNA sequencing from 4,645 cells across 19 patients with metastatic melanoma [9]. We used the authors’ malignant-cell indicator and nonmalignant annotations. Cells with unresolved compartment identity were excluded from localization analyses, leaving 4,097 annotated cells: 1,257 malignant cells, 2,040 T cells, 512 B cells, 119 macrophages, 62 endothelial cells, 56 cancer-associated fibroblasts, and 51 NK cells. Analyses retained patient identity throughout.

### Gene sets and continuous scores

The candidate glutamine-associated set contained 22 genes spanning glutamine handling and connected nitrogen/proline pathways; the candidate IFN-γ set contained 23 immune-response genes adapted from published IFN-γ-related profiles [1,4]. Bulk expression was log2 transformed where needed, and sample scores were calculated with GSVA and standardized within cohort [10]. Sensitivity analyses used MSigDB REACTOME_GLUTAMATE_AND_GLUTAMINE_METABOLISM and HALLMARK_INTERFERON_GAMMA_RESPONSE [11]. Gene mapping was documented for every cohort.

### Bulk composition analyses

We first estimated the Pearson correlation between continuous candidate scores in TCGA. We then residualized each score for (i) ABSOLUTE purity and sample type, (ii) purity, sample type, and EPIC-derived immune/stromal fractions, and (iii) purity, sample type, and principal components of MCP-counter immune/stromal abundance scores [12–14]. EPIC fits with nonconvergence codes were excluded from EPIC-based models (20 of 469 samples). Correlations between each transcriptional score and deconvolution summaries were also calculated. EPIC- and MCP-counter-adjusted Cox models tested whether survival estimates persisted after composition adjustment.

### Single-cell localization

Within GSE72056, each mapped signature gene was standardized across cells; the mean standardized expression of mapped genes defined a cell-level score. Scores were then summarized as the median within each patient-by-compartment combination. Malignant-cell scores were compared with each paired nonmalignant compartment using two-sided paired Wilcoxon tests with Holm adjustment across compartments. Median paired differences were accompanied by bootstrap 95% confidence intervals. Patient-level Spearman correlations assessed malignant glutamine-associated scores against malignant or aggregated immune IFN-γ-associated scores. The patient—not the cell—was the independent unit, avoiding pseudoreplication [20].

### Outcome analyses

Cox proportional-hazards models used one-standard-deviation increments. TCGA models adjusted for age, sex, and grouped stage; GSE65904 models adjusted for age, sex, stage, and tissue source. Candidate models included both score main effects and their product interaction. Nested likelihood-ratio tests evaluated whether IFN-γ, glutamine, and the interaction added fit sequentially to clinical covariates. Proportional-hazards assumptions were assessed with scaled Schoenfeld residuals. Curated signatures and composition-adjusted models were prespecified sensitivity analyses.

GSE91061 response associations used Firth penalized logistic regression because only 10 responses were observed [17]. Leave-one-out cross-validated predictions were summarized by ROC AUC with bootstrap confidence intervals [18]. Monte Carlo simulations used the observed sample size and response prevalence to estimate power across true odds ratios; this analysis quantifies the large effects detectable by the dataset rather than retroactively proving equivalence.

### Missingness, multiplicity, and software

We reported cohort flow and compared score distributions and observed covariates between included and excluded cases. Primary hypotheses were estimated with continuous scores. Single-cell compartment comparisons used Holm adjustment; other sensitivity analyses are identified as such and interpreted by effect magnitude, confidence intervals, and consistency rather than isolated p-value thresholds. Analyses used R 4.4.2 with survival, GSVA, GEOquery, EPIC, MCPcounter, logistf, pROC, and related packages. Exact versions are recorded in the supplied session-information files.

## Results

### Cohorts and score mapping

The four datasets contributed complementary evidence (Table 1 and Figure 1). Candidate signatures mapped nearly completely in every dataset: 22 glutamine and 22 IFN-γ genes in TCGA, 21 and 23 in GSE65904, and 21 and 23 in the single-cell dataset. Curated sets mapped less completely for the small glutamine set but broadly for the 200-gene IFN-γ set. Cohort-specific mapping is reported in Supplementary Tables S2 and S5.

**Figure 1.**
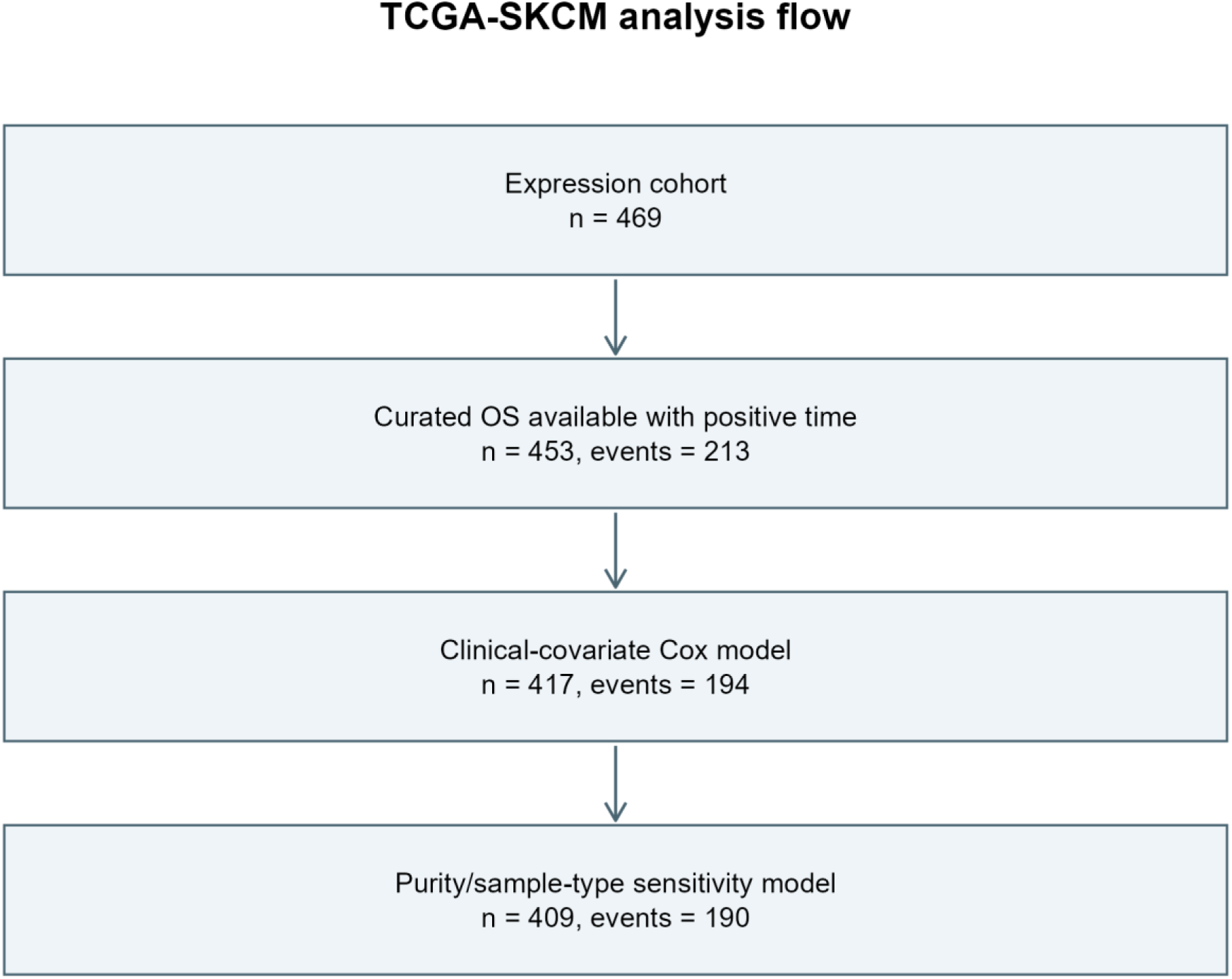
TCGA-SKCM analysis flow from aligned expression profiles to the overall-survival, complete-covariate, and composition-adjusted analysis sets. Table 1 reports the complementary external cohorts.

**Table 1.**
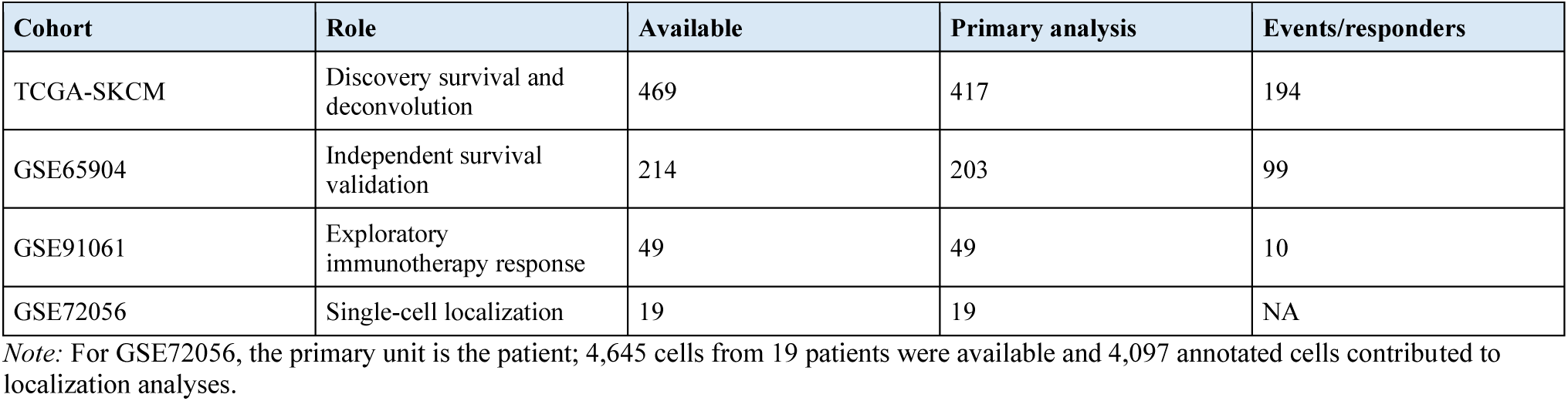
Cohort roles and primary analysis sets.

**Table 2.**
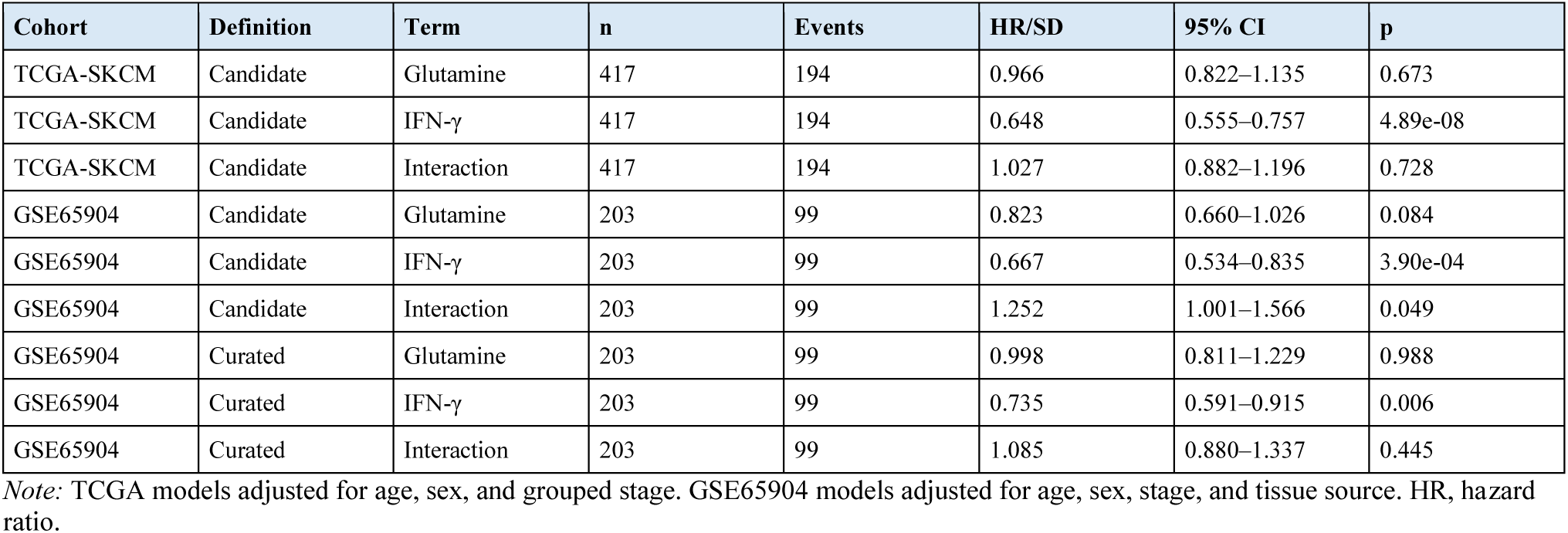
Cross-cohort multivariable survival associations for continuous scores.

### The bulk inverse association is strongly composition dependent

In TCGA, candidate glutamine-associated and IFN-γ-associated scores were inversely correlated (r=−0.397; 95% CI −0.470 to −0.318; p=3.89×10⁻¹⁹; Figure 2). Adjustment for purity and sample type attenuated the association to r=−0.218. Adding EPIC immune/stromal fractions attenuated it further to r=−0.146, whereas adjustment using MCP-counter immune/stromal principal components produced essentially no residual association (r=0.019; 95% CI −0.072 to 0.110; p=0.681; Table 3 and Figure 4).

**Figure 2.**
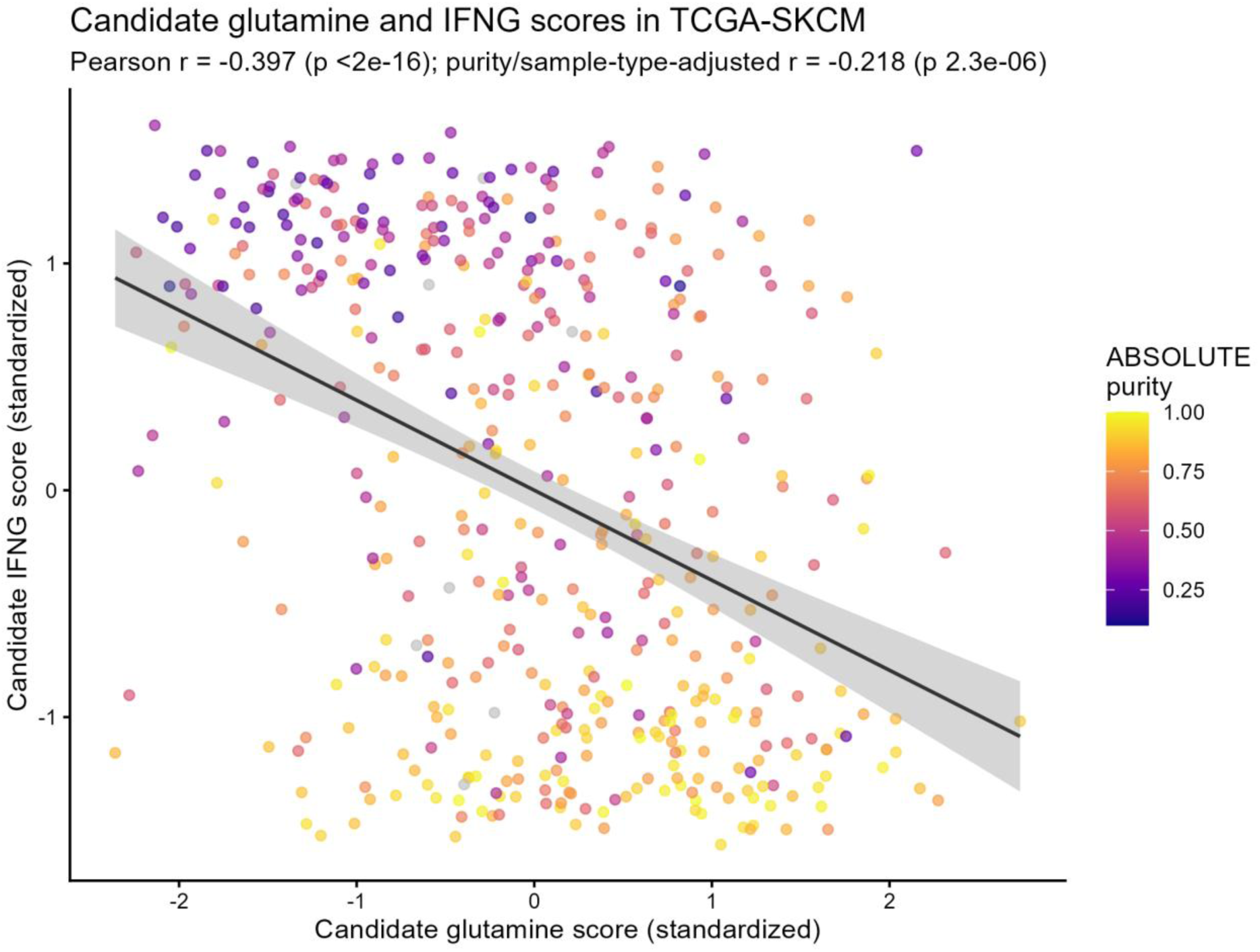
Inverse association between continuous candidate glutamine-associated and IFN-γ-associated GSVA scores in TCGA-SKCM. The line is the least-squares fit; the shaded band is its 95% confidence interval. This unadjusted bulk association is not interpreted as cell-intrinsic antagonism.

**Figure 3.**
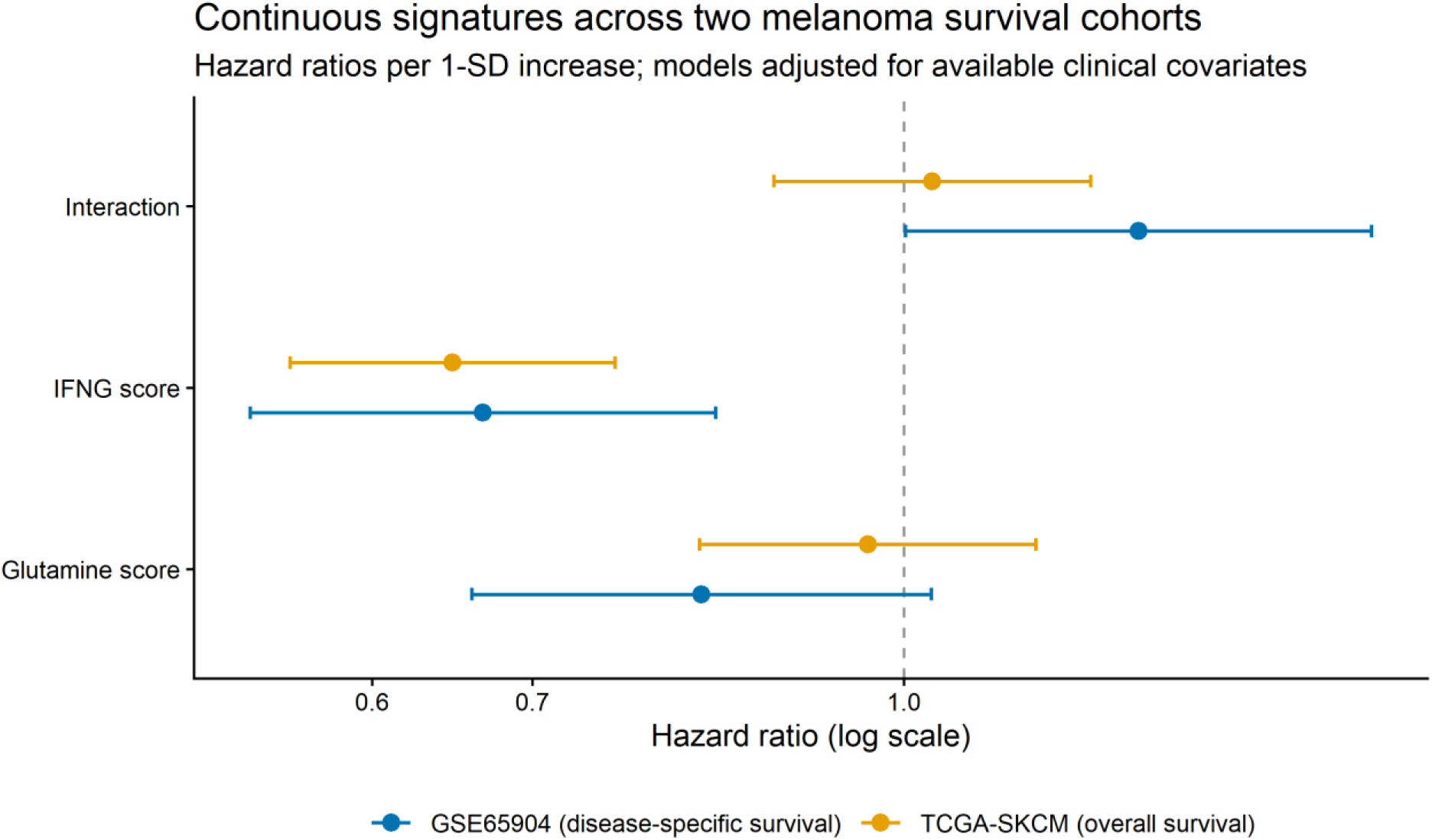
Cross-cohort adjusted survival associations per one-standard-deviation increase in score. IFN-γ-associated transcription is directionally consistent and statistically supported in TCGA OS and GSE65904 DSS. Glutamine and interaction estimates are not consistently supported. The dashed line marks HR=1.

**Figure 4.**
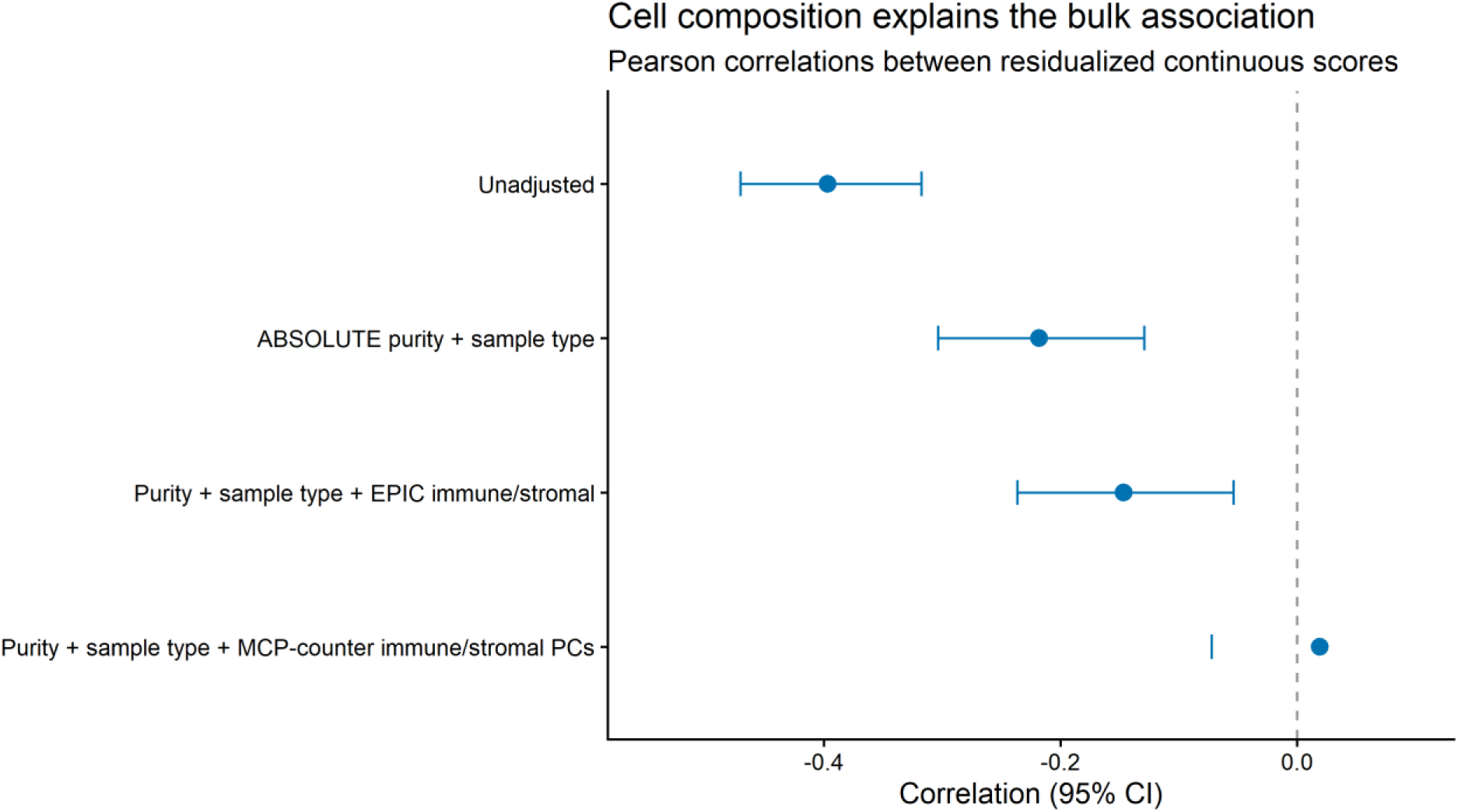
Progressive attenuation of the TCGA bulk glutamine–IFN-γ score correlation after adjustment for tumor purity, sample type, and immune/stromal composition. Points show Pearson correlations of residualized continuous scores; bars show 95% confidence intervals.

**Table 3.**
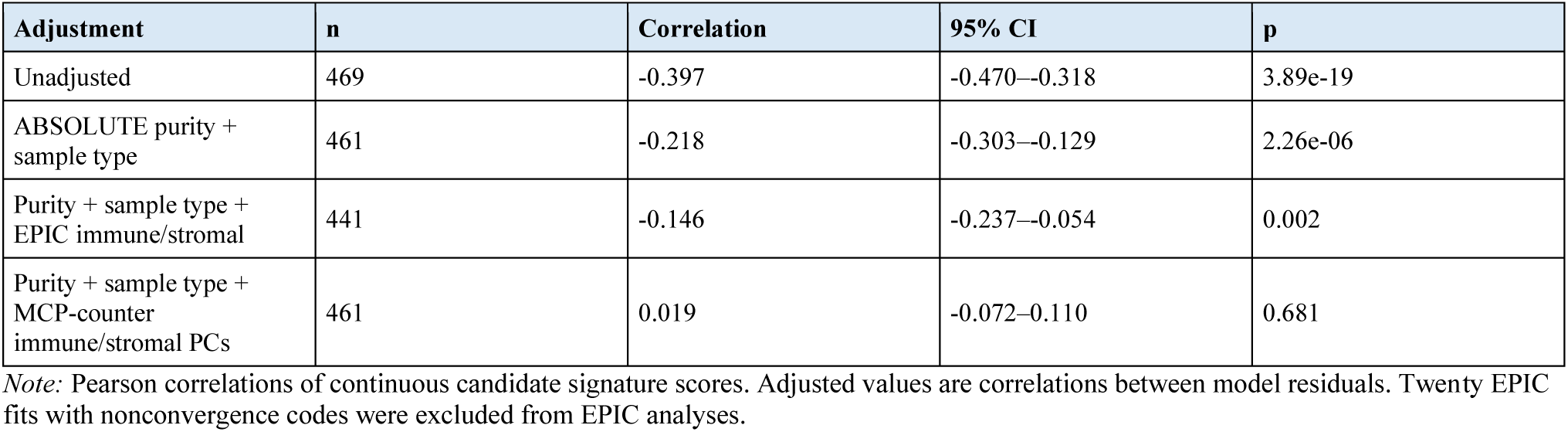
Attenuation of the TCGA bulk score correlation after composition adjustment.

**Table 4.**
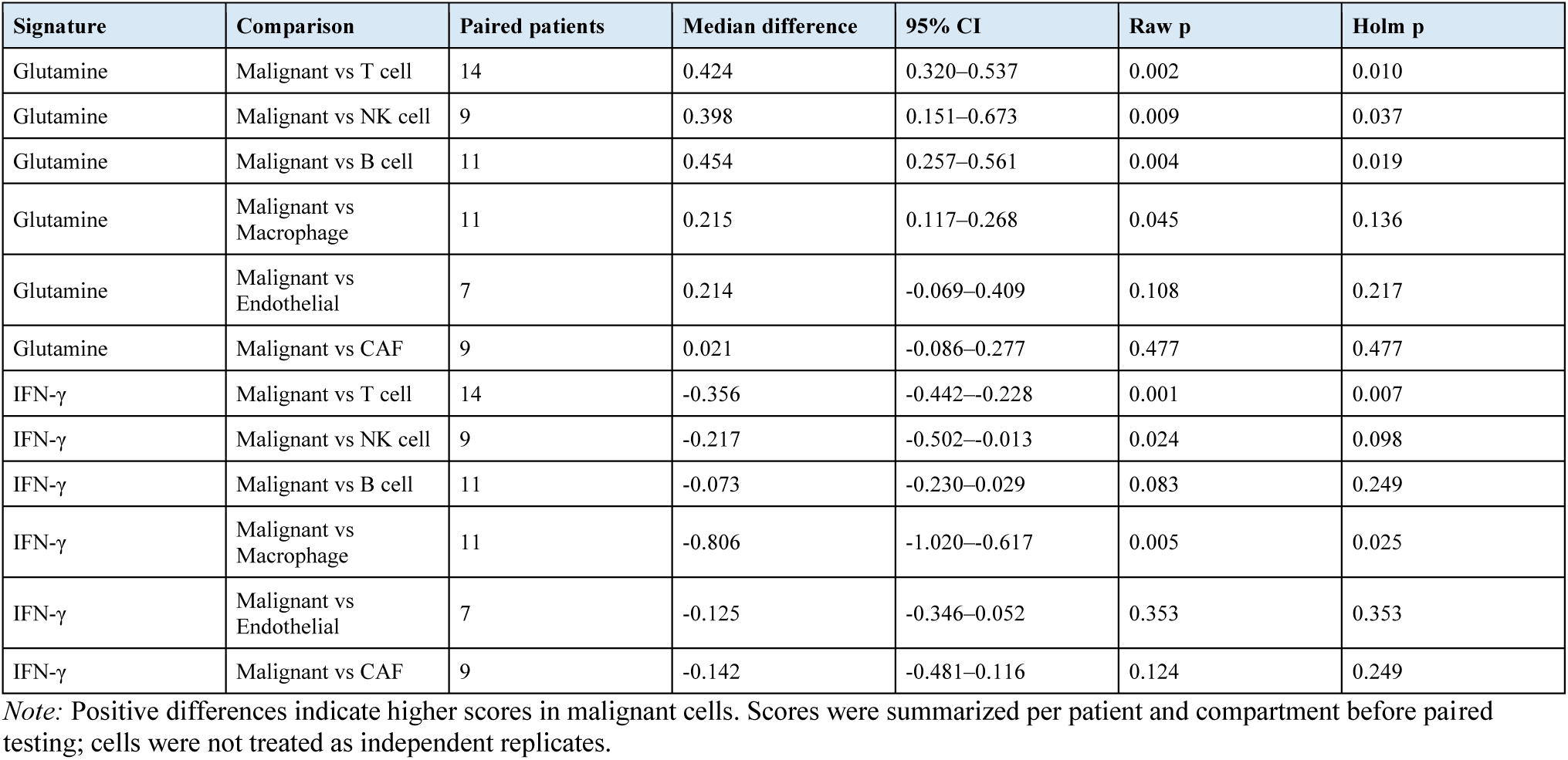
Patient-level single-cell localization of candidate signatures.

**Table 5.**
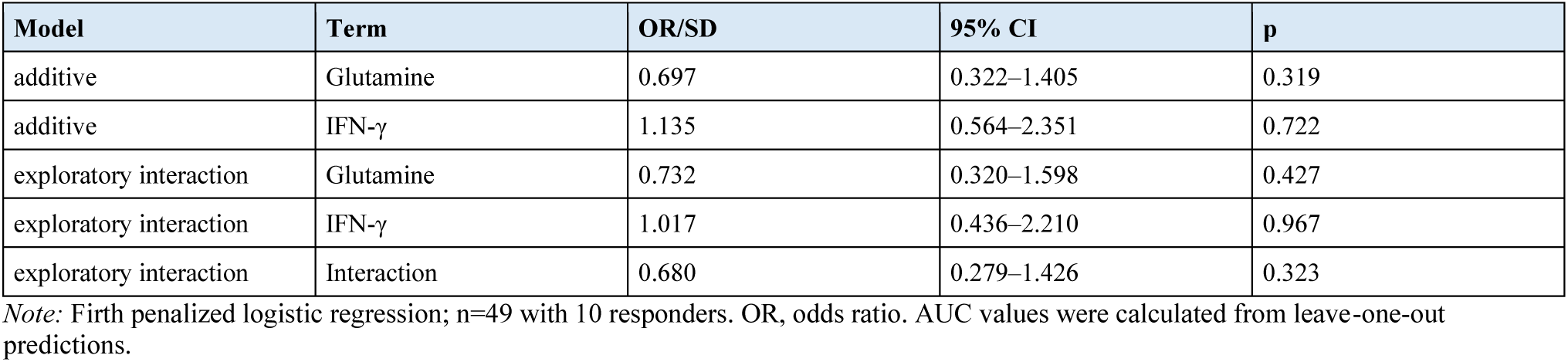
Exploratory pretreatment response models in GSE91061.

The candidate glutamine score was negatively correlated with inferred immune abundance by both EPIC (r=−0.448) and MCP-counter (r=−0.481), while the IFN-γ score was positively correlated with immune abundance (r=0.520 and r=0.800, respectively). This opposing compartment loading explains why the unadjusted bulk scores appear strongly antagonistic. The difference between EPIC and MCP-counter residual estimates also shows that the exact adjusted value depends on the composition model; the robust conclusion is substantial attenuation, not a single corrected correlation.

### Single-cell analysis localizes the programs to different compartments

Patient-level single-cell summaries supported a compartmental explanation (Figure 5). Candidate glutamine-associated scores were higher in malignant cells than paired T cells (median difference 0.424; Holm p=0.010), NK cells (0.398; p=0.037), or B cells (0.454; p=0.019). Candidate IFN-γ-associated scores were lower in malignant cells than paired T cells (median difference −0.356; Holm p=0.0066) and macrophages (−0.806; p=0.026). Curated signatures yielded directionally similar localization, although significance varied with mapping and the small number of paired patients.

**Figure 5.**
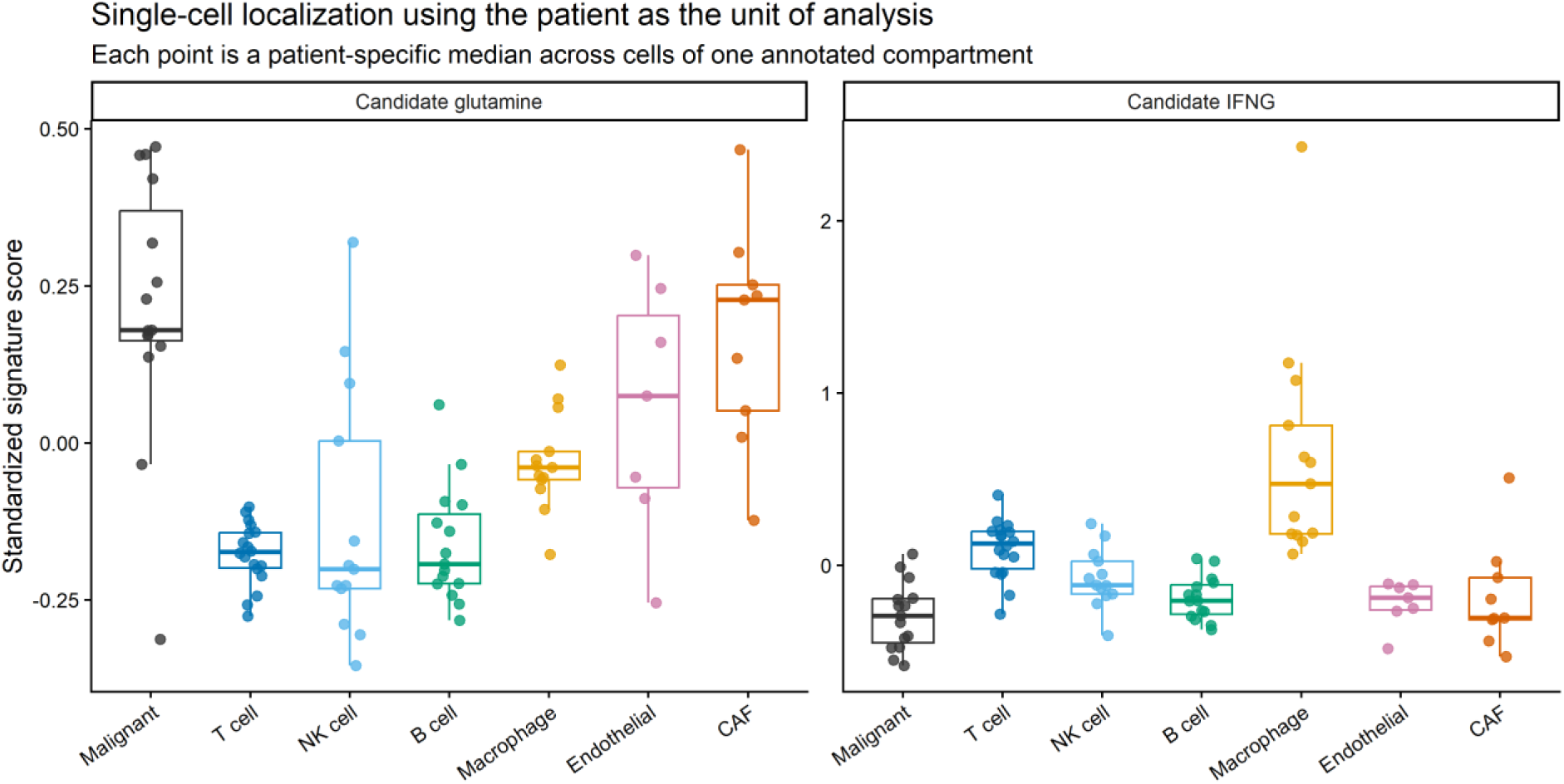
Patient-level single-cell localization in GSE72056. Each point is a patient-specific median score for one annotated compartment; boxplots summarize patients, not cells. Candidate glutamine-associated scores are relatively enriched in malignant cells, whereas IFN-γ-associated scores are enriched in immune compartments.

Across 15 evaluable patients, the correlation between malignant-cell candidate glutamine scores and aggregated immune-cell candidate IFN-γ scores was imprecise (Spearman ρ=−0.386; bootstrap 95% CI −0.842 to 0.256; p=0.156). Thus, cell-type localization is supported, but this small dataset does not establish a patient-level antagonistic mechanism.

### IFN-γ prognosis replicates; glutamine prognosis does not

In the TCGA primary multivariable model (n=417; 194 deaths), higher IFN-γ-associated score was associated with lower mortality (HR per SD 0.648; 95% CI 0.555–0.757; p=4.89×10⁻⁸), whereas the glutamine-associated score was not (HR 0.966; 95% CI 0.822–1.135; p=0.673). Adding IFN-γ to the clinical model improved fit (likelihood-ratio p=8.20×10⁻⁹; concordance 0.636 to 0.691). Adding glutamine after IFN-γ did not improve fit (p=0.616).

The IFN-γ association replicated in GSE65904 (n=203; 99 DSS deaths; adjusted HR 0.667; 95% CI 0.534–0.835; p=3.90×10⁻⁴). The glutamine estimate was directionally protective but uncertain (HR 0.823; 95% CI 0.660–1.026; p=0.084). IFN-γ also improved the external clinical model (likelihood-ratio p=0.000505; concordance 0.621 to 0.676). Curated signatures confirmed the IFN-γ result in both cohorts and did not support an independent glutamine association. Figure 3 summarizes cross-cohort estimates.

Composition-adjusted TCGA models did not materially change the IFN-γ result: HR 0.668 (95% CI 0.550–0.811; p=4.40×10⁻⁵) after EPIC adjustment and HR 0.774 (95% CI 0.596–1.004; p=0.054) after MCP-counter adjustment. The latter estimate was less precise but directionally consistent. Glutamine remained null. Proportional-hazards tests did not identify a global violation in the external model.

### The interaction is not a replicated biomarker

The candidate glutamine-by-IFN-γ interaction was null in TCGA (HR 1.027; 95% CI 0.882–1.196; p=0.728). In GSE65904, the candidate interaction was nominally associated with DSS (HR 1.252; 95% CI 1.001–1.566; p=0.049), and the nested interaction test gave p=0.044. However, the curated-signature interaction was null in GSE65904 (HR 1.085; p=0.445), and neither candidate nor curated interactions were supported in TCGA. We therefore classify the external candidate interaction as an unstable, exploratory observation rather than validation.

### The treated cohort is too small for modest response effects

In GSE91061, neither candidate score was associated with response in Firth models: glutamine odds ratio (OR) 0.697 (95% CI 0.322–1.405; p=0.319) and IFN-γ OR 1.135 (95% CI 0.564–2.351; p=0.722). The exploratory interaction OR was 0.680 (95% CI 0.279–1.426; p=0.323). Leave-one-out AUC was 0.356 (95% bootstrap CI 0.156–0.556; Figure 6). All score intervals were compatible with both decreases and increases in response odds.

**Figure 6.**
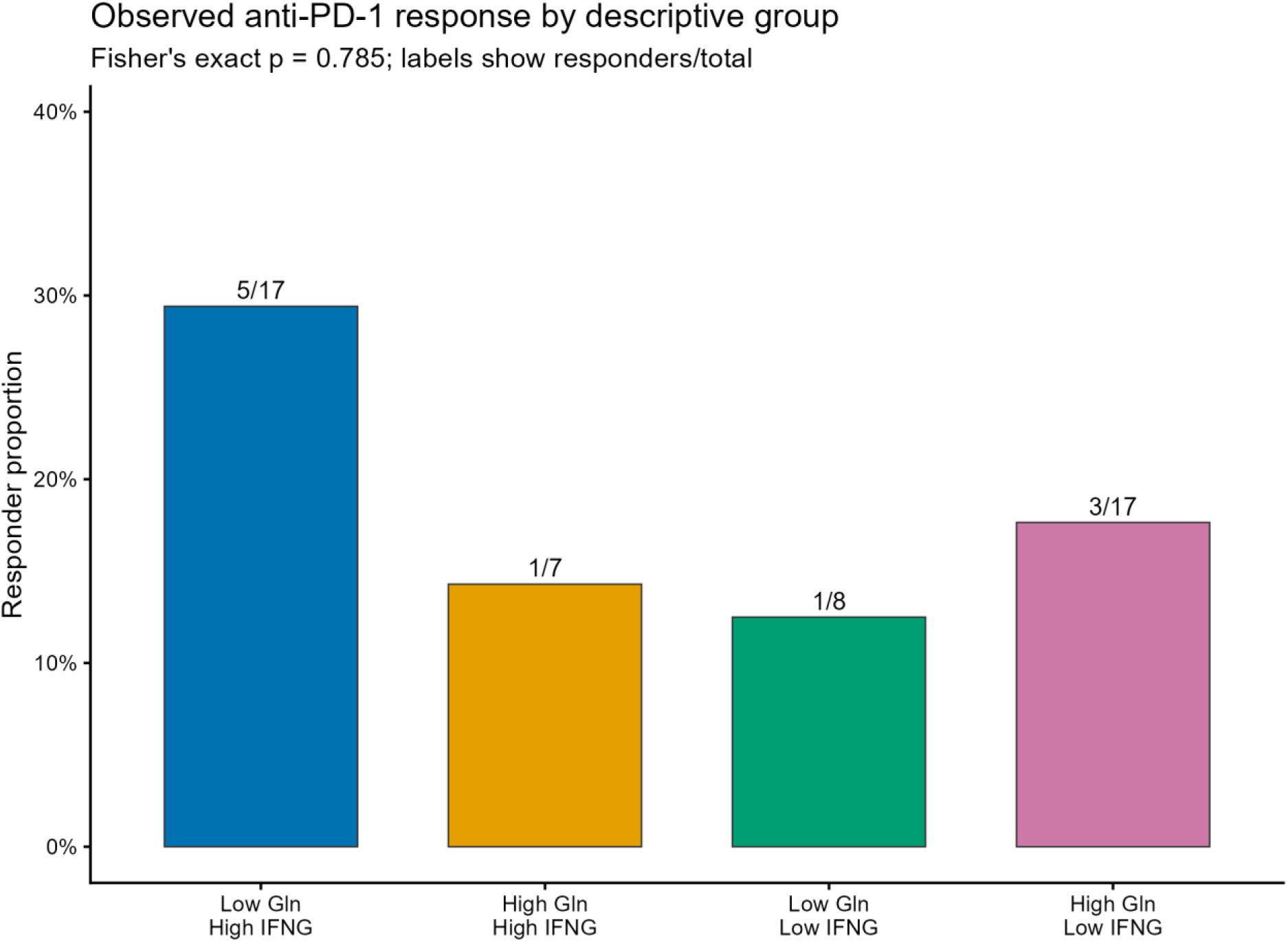
Exploratory pretreatment score distributions by response in GSE91061. The cohort contained 49 tumors and 10 responders. Firth regression and leave-one-out discrimination did not support a response association, but confidence intervals were wide.

At the observed prevalence of 10 responses among 49 patients, simulation indicated that an OR of approximately 3.25 per SD was required to reach 80% power under the modeled conditions (Supplementary Figure S3). Accordingly, the null estimates do not exclude modest associations. They do show that this cohort cannot credibly validate a multivariable predictive signature.

### Missingness and robustness

TCGA patients included in the complete-covariate model had score distributions similar to patients with expression but incomplete outcome/covariate data. Stage distribution differed between OS-eligible and expression-only groups (p=0.035), and stage missingness drove most primary-model exclusions. Included patients had shorter observed OS time than OS-eligible excluded patients, while score distributions remained similar. These patterns limit transportability and reinforce interpretation of the models as adjusted associations, not population-level risk tools.

## Discussion

### Principal findings

This strengthened analysis changes the biological interpretation of the original observation. The inverse relationship between glutamine-associated and IFN-γ-associated transcription is real in bulk melanoma RNA, but it is not robust to modeling cellular composition. Independent deconvolution methods and single-cell localization converge on a parsimonious explanation: glutamine-associated transcription is relatively enriched in malignant cells, whereas IFN-γ-associated transcription is concentrated in immune compartments. Variation in the mixture of these compartments generates much of the apparent bulk antagonism.

The most reproducible outcome result is not a two-score metabolic-conflict biomarker but the favorable prognostic association of IFN-γ-associated transcription. This association replicated across OS and DSS cohorts, survived curated-signature analyses, and remained directionally consistent after composition adjustment. Glutamine-associated transcription neither consistently predicted survival nor improved models after IFN-γ. The nominal external interaction did not reproduce across signatures or cohorts and should not motivate clinical stratification without new data.

### Biological interpretation

The findings do not refute metabolic competition. Experimental work shows that nutrient restriction can alter tumor and immune-cell function [2,3]. Rather, they demonstrate that bulk coexpression is insufficient evidence for that mechanism. A score can reflect cellular abundance, a cell-intrinsic state, or both. Here, the near-null MCP-counter-adjusted correlation and the single-cell compartment differences place cellular mixture at the center of the bulk signal.

The remaining uncertainty is mechanistic and spatial. The single-cell dataset had only 19 patients, lacked matched functional metabolite measurements, and used dissociated cells rather than spatially resolved neighborhoods. Patient-level cross-compartment correlations had wide confidence intervals. Future work should pair spatial transcriptomics or multiplex imaging with isotope tracing, nutrient measurements, or perturbation assays to determine whether glutamine-active malignant neighborhoods suppress nearby IFN-γ programs.

### Clinical interpretation

IFN-γ-associated transcription is prognostic in these datasets, but prognostic association does not establish immunotherapy prediction. Demonstrating predictive value requires treatment-by-biomarker interaction in a randomized or otherwise credible treatment-comparative design. The single treated cohort analyzed here is small, includes only 10 responders, and can detect only very large effects. The present data therefore provide no basis for treatment selection or clinical cut points.

### Strengths and limitations

Strengths include independent survival validation, continuous-score modeling, curated-signature sensitivity analyses, three complementary composition approaches, patient-level single-cell inference, explicit incremental-value tests, small-sample response methods, and complete machine-readable outputs. These elements address the principal weaknesses of the preliminary version.

Limitations remain. All analyses are retrospective. Cohorts differed in platform, tissue source, endpoint, and clinical annotation. Deconvolution relies on reference profiles and model assumptions; EPIC did not fully converge for 20 tumors, which were excluded from EPIC models. The single-cell data include few patients and uneven cell-type recovery. Signature scores are transcriptional proxies rather than glutamine flux or IFN-γ protein activity. Missing clinical covariates may induce selection bias. Multiple sensitivity analyses increase the chance of nominal findings, particularly the isolated GSE65904 interaction. Finally, no treatment-comparative cohort was available.

## Conclusions

Cellular composition accounts for much of the inverse bulk glutamine–IFN-γ transcriptional axis in melanoma. IFN-γ-associated transcription is a reproducible favorable prognostic correlate, but glutamine-associated transcription and the cross-score interaction do not provide stable incremental outcome information. The results support cell-resolved mechanistic studies and adequately powered treatment-comparative validation—not a median-defined two-score clinical classifier.

## Supporting information

Editable Manuscript Source File

## Declarations

### Ethics approval and consent

Only public, deidentified data were analyzed; no new participant recruitment or intervention was performed. Ethical approvals and consent procedures are described in the source studies.

### Data availability

TCGA-SKCM data are available through the NCI Genomic Data Commons. GEO datasets are available under GSE65904, GSE91061, and GSE72056. The preliminary preprint is available at doi: 10.1101/2025.09.28.679008.

### Code availability

Complete analysis scripts, derived tables, figures, data-acquisition manifests, and software session information are available in GitHub release v2.0.1 at https://github.com/DataStalker/Melanoma-Metabolic-Conflict-Analysis/releases/tag/v2.0.1 and are permanently archived on Zenodo at https://doi.org/10.5281/zenodo.21895340.

### Competing interests

Islam Asal is the founder of OncoMetrika. The author declares no other competing interests.

### Funding

No external funding was received for this work.

### Author contributions

Islam Asal: conceptualization, methodology, software, formal analysis, visualization, writing—original draft, and writing—review and editing.

## Acknowledgments

The author thanks the investigators and participants who generated and shared the TCGA and GEO datasets, and the PREreview contributors whose methodological critique motivated the strengthened analysis.

## Notes

https://www.ncbi.nlm.nih.gov/geo/query/acc.cgi?acc=GSE91061

https://portal.gdc.cancer.gov/projects/TCGA-SKCM

https://www.ncbi.nlm.nih.gov/geo/query/acc.cgi?acc=GSE65904

https://www.ncbi.nlm.nih.gov/geo/query/acc.cgi?acc=GSE72056

https://github.com/DataStalker/Melanoma-Metabolic-Conflict-Analysis/releases/tag/v2.0.1

https://doi.org/10.5281/zenodo.21895340

### Summary of Updates

This revision corrects endpoint harmonization and expands the original bulk transcriptomic analysis. It adds independent disease specific survival validation in GSE65904, tumor purity and cell composition adjustment using ABSOLUTE, EPIC, and MCP-counter, patient level single cell localization in GSE72056, curated signature sensitivity analyses, missingness diagnostics, and response effect precision and power analyses. The conclusions have been revised: cellular composition accounts for much of the bulk glutamine and IFN-gamma relationship; IFN-gamma associated transcription is prognostic across two cohorts, whereas glutamine associated transcription and the interaction do not provide stable incremental outcome information. Code and derived results are now archived on GitHub and Zenodo.

## References

1. Ratnikov B, Aza-Blanc P, Ronai ZA, Smith JW, Osterman AL, Scott DA. Glutamate and asparagine cataplerosis underlie glutamine addiction in melanoma. Oncotarget. 2015;6:7379–7389. doi:10.18632/oncotarget.3132.

2. Chang CH, Qiu J, O’Sullivan D, et al. Metabolic competition in the tumor microenvironment is a driver of cancer progression. Cell. 2015;162:1229–1241. doi:10.1016/j.cell.2015.08.016.

3. Leone RD, Zhao L, Englert JM, et al. Glutamine blockade induces divergent metabolic programs to overcome tumor immune evasion. Science. 2019;366:1013–1021. doi:10.1126/science.aav2588.

4. Ayers M, Lunceford J, Nebozhyn M, et al. IFN-γ-related mRNA profile predicts clinical response to PD-1 blockade. J Clin Invest. 2017;127:2930–2940. doi:10.1172/JCI91190.

5. Cancer Genome Atlas Network. Genomic classification of cutaneous melanoma. Cell. 2015;161:1681–1696. doi:10.1016/j.cell.2015.05.044.

6. Riaz N, Havel JJ, Makarov V, et al. Tumor and microenvironment evolution during immunotherapy with nivolumab. Cell. 2017;171:934–949.e16. doi:10.1016/j.cell.2017.09.028.

7. Liu J, Lichtenberg T, Hoadley KA, et al. An integrated TCGA pan-cancer clinical data resource to drive high-quality survival outcome analytics. Cell. 2018;173:400–416.e11. doi:10.1016/j.cell.2018.02.052.

8. Cirenajwis H, Ekedahl H, Lauss M, et al. Molecular stratification of metastatic melanoma using gene expression profiling: prediction of survival outcome and benefit from molecular targeted therapy. Oncotarget. 2015;6:12297–12309. doi:10.18632/oncotarget.3655.

9. Tirosh I, Izar B, Prakadan SM, et al. Dissecting the multicellular ecosystem of metastatic melanoma by single-cell RNA-seq. Science. 2016;352:189–196. doi:10.1126/science.aad0501.

10. Hänzelmann S, Castelo R, Guinney J. GSVA: gene set variation analysis for microarray and RNA-seq data. BMC Bioinformatics. 2013;14:7. doi:10.1186/1471-2105-14-7.

11. Liberzon A, Birger C, Thorvaldsdóttir H, et al. The Molecular Signatures Database Hallmark Gene Set Collection. Cell Syst. 2015;1:417–425. doi:10.1016/j.cels.2015.12.004.

12. Carter SL, Cibulskis K, Helman E, et al. Absolute quantification of somatic DNA alterations in human cancer. Nat Biotechnol. 2012;30:413–421. doi:10.1038/nbt.2203.

13. Becht E, Giraldo NA, Lacroix L, et al. Estimating the population abundance of tissue-infiltrating immune and stromal cell populations using gene expression. Genome Biol. 2016;17:218. doi:10.1186/s13059-016-1070-5.

14. Racle J, de Jonge K, Baumgaertner P, Speiser DE, Gfeller D. Simultaneous enumeration of cancer and immune cell types from bulk tumor gene expression data. eLife. 2017;6:e26476. doi:10.7554/eLife.26476.

15. Davis S, Meltzer PS. GEOquery: a bridge between the Gene Expression Omnibus and BioConductor. Bioinformatics. 2007;23:1846–1847. doi:10.1093/bioinformatics/btm254.

16. Colaprico A, Silva TC, Olsen C, et al. TCGAbiolinks: an R/Bioconductor package for integrative analysis of TCGA data. Nucleic Acids Res. 2016;44:e71. doi:10.1093/nar/gkv1507.

17. Heinze G, Schemper M. A solution to the problem of separation in logistic regression. Stat Med. 2002;21:2409–2419. doi:10.1002/sim.1047.

18. Robin X, Turck N, Hainard A, et al. pROC: an open-source package for R and S+ to analyze and compare ROC curves. BMC Bioinformatics. 2011;12:77. doi:10.1186/1471-2105-12-77.

19. McShane LM, Altman DG, Sauerbrei W, et al. REporting recommendations for tumor MARKer prognostic studies (REMARK). J Natl Cancer Inst. 2005;97:1180–1184. doi:10.1093/jnci/dji237.

20. Squair JW, Gautier M, Kathe C, et al. Confronting false discoveries in single-cell differential expression. Nat Commun. 2021;12:5692. doi:10.1038/s41467-021-25960-2.

21. Sade -Feldman M, Yizhak K, Bjorgaard SL, et al. Defining T cell states associated with response to checkpoint immunotherapy in melanoma. Cell. 2018;175:998–1013.e20. doi:10.1016/j.cell.2018.10.038.

