## Supplementary Methods, Tables and Figures for "Cellular Composition Accounts for Much of the Bulk Glutamine–IFN-γ Transcriptional Axis in Melanoma: A Cross-Cohort and Single-Cell Analysis"

Islam Asal | OncoMetrika, Cairo, Egypt

### Supplementary material

#### Supplementary methods

Data acquisition was scripted and cached with MD5 checksums. GSE65904 series-matrix and GPL10558 annotation files were retrieved from GEO; GSE72056 revised single-cell data were retrieved from GEO; MCP-counter gene markers were pinned to repository commit b6eac73. The acquisition manifest records local filenames, source URLs, sizes, and checksums. All strengthened analyses can be regenerated by running 05_acquire_strengthening_data.R followed by 04_revised_analysis.R and 06_strengthened_analysis.R, subject to availability of the public source files and required R packages.

#### Supplementary results

Curated-signature analyses reproduced the favorable IFN-γ survival association and did not support a stable glutamine or interaction effect. In GSE65904, curated IFN-γ HR was 0.735 (95% CI 0.591–0.915; p=0.0059), curated glutamine HR was 0.998 (p=0.988), and the interaction HR was 1.085 (p=0.445). In TCGA, corresponding estimates were 0.636 ($p=1.44\times10^{-8}$), 0.947 (p=0.507), and 0.992 (p=0.915).

After EPIC adjustment in TCGA, IFN-γ remained associated with survival (HR 0.668; $p=4.40\times10^{-5}$), glutamine remained null (HR 0.939; p=0.495), and the interaction remained null (HR 0.963; p=0.667). MCP-counter-adjusted estimates were less precise: IFN-γ HR 0.774 (p=0.054), glutamine HR 0.917 (p=0.334), and interaction HR 0.998 (p=0.984).

**Table S1.** Incremental model fit in the independent GSE65904 survival cohort

| Model | n | Events | Parameters | AIC | Concordance | Incremental LRT p |
|---|---|---|---|---|---|---|
| Clinical covariates | 203 | 99 | 6 | 908.79 | 0.621 | — |
| Clinical + IFNG | 203 | 99 | 7 | 898.70 | 0.676 | 5.05e-04 |
| Clinical + IFNG + glutamine | 203 | 99 | 8 | 897.18 | 0.678 | 0.061 |
| Clinical + IFNG + glutamine + interaction | 203 | 99 | 9 | 895.13 | 0.691 | 0.044 |

*Note:* Nested likelihood-ratio tests compare each row with the preceding model. Candidate signatures are shown.

**Table S2.** Signature mapping across cohorts

| Cohort | Signature | Genes specified | Genes mapped |
|---|---|---|---|
| GSE65904 | candidate_glutamine | 22 | 21 |
| GSE65904 | candidate_ifng | 23 | 23 |
| GSE65904 | reactome_glutamine | 14 | 12 |
| GSE65904 | hallmark_ifng | 200 | 193 |
| GSE72056 | candidate_glutamine | 22 | 21 |
| GSE72056 | candidate_ifng | 23 | 23 |
| GSE72056 | reactome_glutamine | 14 | 12 |
| GSE72056 | hallmark_ifng | 200 | 195 |

*Note:* The complete machine-readable mapping tables are supplied with the analysis package.

**Table S3.** TCGA score and covariate missingness comparisons

| Comparison | Variable | Included/eligible | Excluded/other | p |
|---|---|---|---|---|
| OS-eligible vs expression-only | gln_z | 0.03 [-0.75, 0.77] | 0.01 [-0.26, 0.55] | 0.893 |
| OS-eligible vs expression-only | ifng_z | 0.05 [-1.01, 1.00] | -0.51 [-0.84, 0.32] | 0.240 |
| OS-eligible vs expression-only | age | 58.00 [48.00, 71.00] | 55.00 [46.50, 61.75] | 0.592 |
| OS-eligible vs expression-only | purity | 0.70 [0.49, 0.85] | 0.58 [0.45, 0.73] | 0.171 |
| OS-eligible vs expression-only | gender | Female: 173 (38.2%); Male: 280 (61.8%) | Female: 7 (43.8%); Male: 9 (56.2%) | 0.795 |
| OS-eligible vs expression-only | stage | Early (0/I/II): 226 (49.9%); Late (III/IV): 191 (42.2%); NA: 36 (7.9%) | Early (0/I/II): 12 (75.0%); Late (III/IV): 2 (12.5%); NA: 2 (12.5%) | 0.035 |
| OS-eligible vs expression-only | sample_type | Primary: 105 (23.2%); Metastatic: 342 (75.5%); NA: 6 (1.3%) | Primary: 1 (6.2%); Metastatic: 15 (93.8%); NA: 0 (0.0%) | 0.296 |
| Primary multivariable vs OS-only | gln_z | 0.03 [-0.74, 0.77] | 0.01 [-1.19, 0.73] | 0.519 |
| Primary multivariable vs OS-only | ifng_z | 0.03 [-1.00, 0.97] | 0.48 [-1.05, 1.12] | 0.427 |
| Primary multivariable vs OS-only | age | 59.00 [48.00, 71.00] | 51.00 [45.75, 67.50] | 0.251 |
| Primary multivariable vs OS-only | purity | 0.70 [0.49, 0.85] | 0.69 [0.54, 0.85] | 0.840 |
| Primary multivariable vs OS-only | os_time | 1035.00 [478.00, 2193.00] | 1546.00 [950.50, 4318.00] | 0.025 |
| Primary multivariable vs OS-only | gender | Female: 158 (37.9%); Male: 259 (62.1%) | Female: 15 (41.7%); Male: 21 (58.3%) | 0.721 |
| Primary multivariable vs OS-only | stage | Early (0/I/II): 226 (54.2%); Late (III/IV): 191 (45.8%); NA: 0 (0.0%) | Early (0/I/II): 0 (0.0%); Late (III/IV): 0 (0.0%); NA: 36 (100.0%) | 1.00e-04 |
| Primary multivariable vs OS-only | sample_type | Primary: 99 (23.7%); Metastatic: 312 (74.8%); NA: 6 (1.4%) | Primary: 6 (16.7%); Metastatic: 30 (83.3%); NA: 0 (0.0%) | 0.637 |
| Primary multivariable vs OS-only | os_event | 0: 223 (53.5%); 1: 194 (46.5%) | 0: 17 (47.2%); 1: 19 (52.8%) | 0.491 |

*Note:* Summaries and tests are exploratory diagnostics of selection into analytic sets.

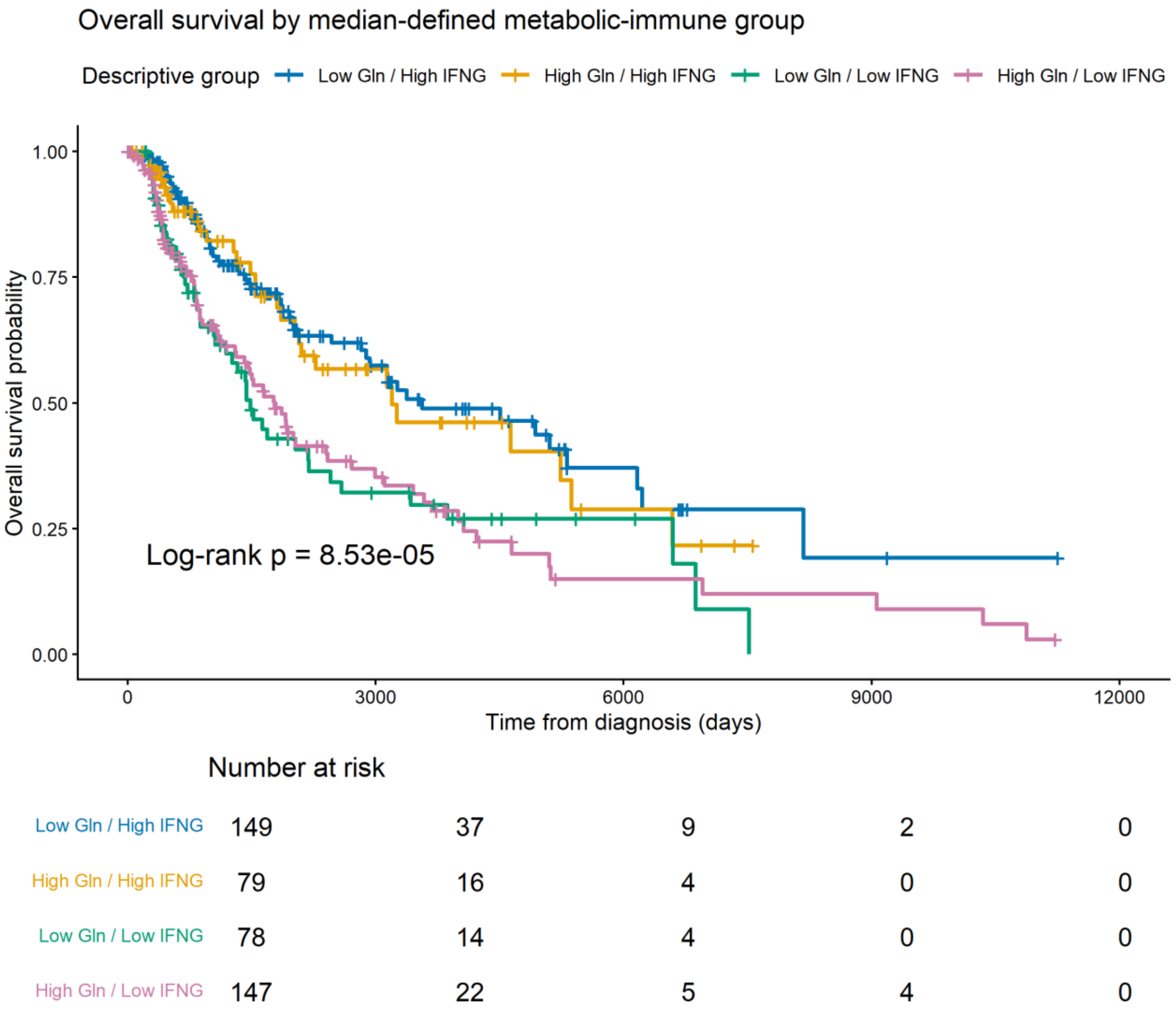


**Figure S1.** Descriptive TCGA Kaplan–Meier curves for median-defined score groups. These groups were not the primary estimand and should not be used as clinical cut points.

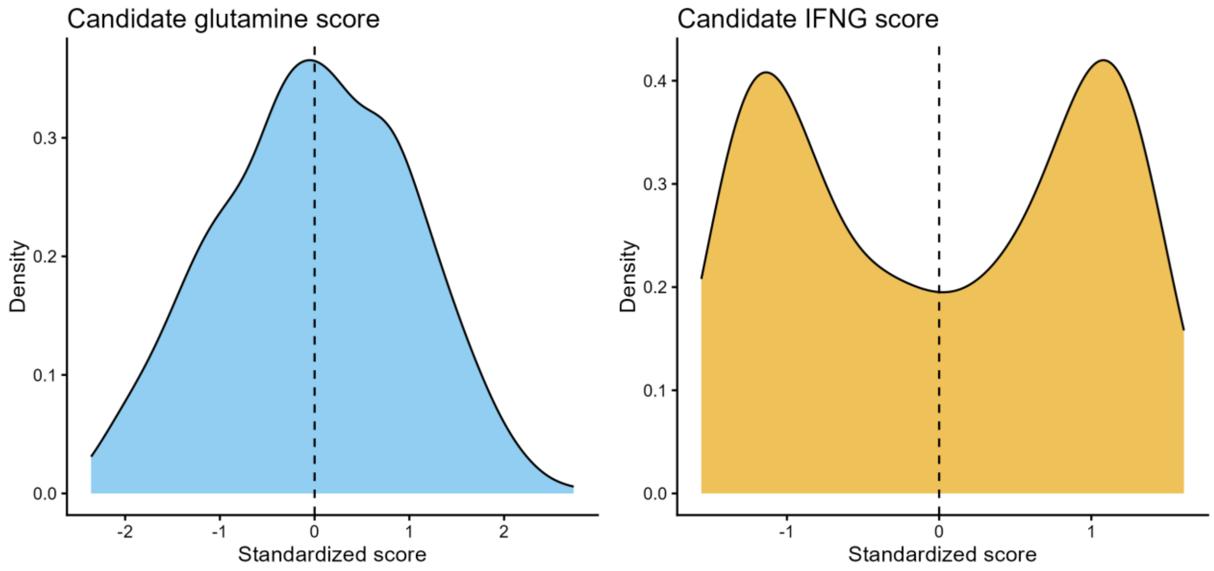


**Figure S2.** Continuous score distributions in TCGA-SKCM and GSE91061.

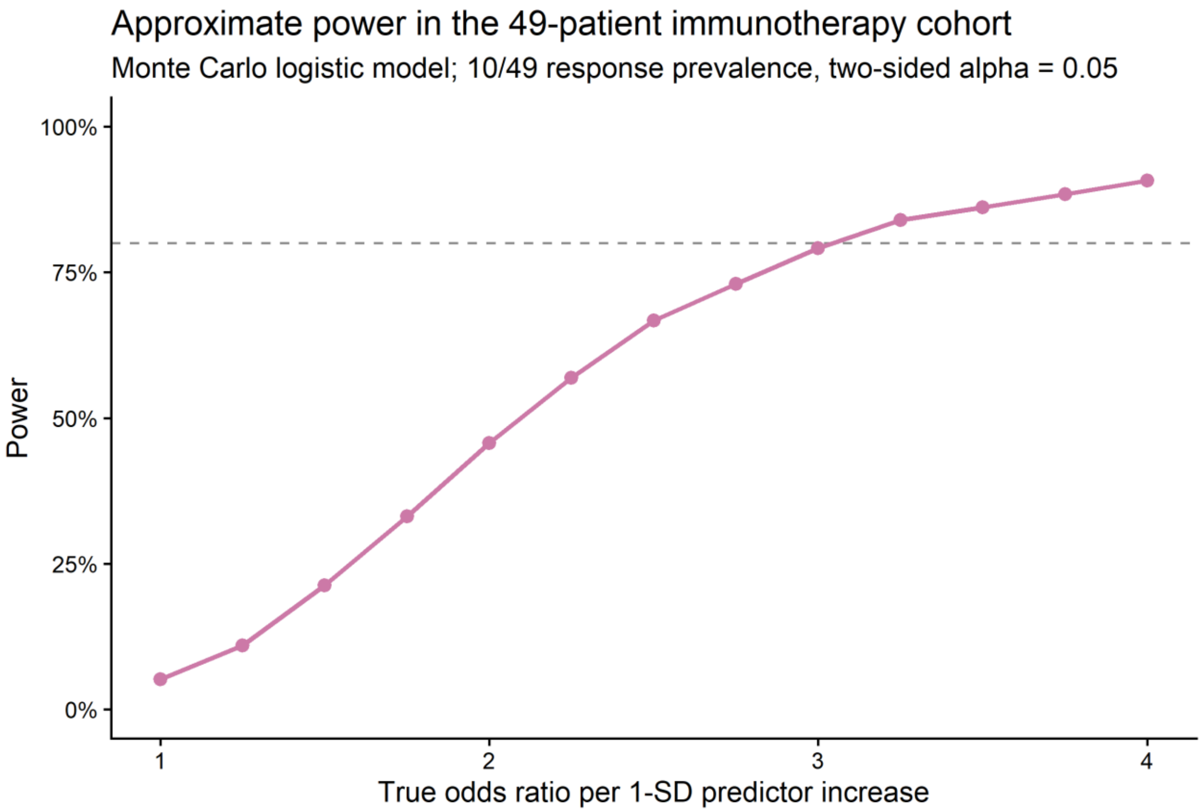


**Figure S3.** Monte Carlo power under the observed GSE91061 sample size (n=49) and response prevalence (10/49). Approximately OR=3.25 per SD was required for 80% power under the simulation model.
