## Supplementary material for "Cellular Composition Accounts for Much of the Bulk Glutamine–IFN-γ Transcriptional Axis in Melanoma: A Cross-Cohort and Single-Cell Analysis": Editable Manuscript Source File

**Islam Asal**

OncoMetrika, Cairo, Egypt

ORCID: 0009-0004-3187-7945

**Article type:** Original research / transcriptomic biomarker evaluation **Running title:** Cellular composition of the glutamine–IFN-γ axis **Keywords:** melanoma; glutamine metabolism; interferon gamma; single-cell RNA sequencing; tumor microenvironment; survival; immunotherapy **Preprint:** A preliminary version was posted on bioRxiv (doi: 10.1101/2025.09.28.679008). This manuscript reports a corrected and substantially expanded analysis.

**Central finding**
The inverse glutamine–IFN-γ relationship in bulk melanoma RNA is largely explained by cellular composition. IFN-γ-associated transcription is independently prognostic in two cohorts, whereas glutamine-associated transcription and the cross-score interaction are not consistently informative.

### Tables

**Table 1.** Cohort roles and primary analysis sets

| **Cohort** | **Role** | **Available** | **Primary analysis** | **Events/responders** |
| --- | --- | --- | --- | --- |
| TCGA-SKCM | Discovery survival and deconvolution | 469 | 417 | 194 |
| GSE65904 | Independent survival validation | 214 | 203 | 99 |
| GSE91061 | Exploratory immunotherapy response | 49 | 49 | 10 |
| GSE72056 | Single-cell localization | 19 | 19 | NA |

*Note:* For GSE72056, the primary unit is the patient; 4,645 cells from 19 patients were available and 4,097 annotated cells contributed to localization analyses.

**Table 2.** Cross-cohort multivariable survival associations for continuous scores

| **Cohort** | **Definition** | **Term** | **n** | **Events** | **HR/SD** | **95% CI** | **p** |
| --- | --- | --- | --- | --- | --- | --- | --- |
| TCGA-SKCM | Candidate | Glutamine | 417 | 194 | 0.966 | 0.822–1.135 | 0.673 |
| TCGA-SKCM | Candidate | IFN-γ | 417 | 194 | 0.648 | 0.555–0.757 | 4.89e-08 |
| TCGA-SKCM | Candidate | Interaction | 417 | 194 | 1.027 | 0.882–1.196 | 0.728 |
| GSE65904 | Candidate | Glutamine | 203 | 99 | 0.823 | 0.660–1.026 | 0.084 |
| GSE65904 | Candidate | IFN-γ | 203 | 99 | 0.667 | 0.534–0.835 | 3.90e-04 |
| GSE65904 | Candidate | Interaction | 203 | 99 | 1.252 | 1.001–1.566 | 0.049 |
| GSE65904 | Curated | Glutamine | 203 | 99 | 0.998 | 0.811–1.229 | 0.988 |
| GSE65904 | Curated | IFN-γ | 203 | 99 | 0.735 | 0.591–0.915 | 0.006 |
| GSE65904 | Curated | Interaction | 203 | 99 | 1.085 | 0.880–1.337 | 0.445 |

*Note:* TCGA models adjusted for age, sex, and grouped stage. GSE65904 models adjusted for age, sex, stage, and tissue source. HR, hazard ratio.

**Table 3.** Attenuation of the TCGA bulk score correlation after composition adjustment

| **Adjustment** | **n** | **Correlation** | **95% CI** | **p** |
| --- | --- | --- | --- | --- |
| Unadjusted | 469 | -0.397 | -0.470–-0.318 | 3.89e-19 |
| ABSOLUTE purity + sample type | 461 | -0.218 | -0.303–-0.129 | 2.26e-06 |
| Purity + sample type + EPIC immune/stromal | 441 | -0.146 | -0.237–-0.054 | 0.002 |
| Purity + sample type + MCP-counter immune/stromal PCs | 461 | 0.019 | -0.072–0.110 | 0.681 |

*Note:* Pearson correlations of continuous candidate signature scores. Adjusted values are correlations between model residuals. Twenty EPIC fits with nonconvergence codes were excluded from EPIC analyses.

**Table 4.** Patient-level single-cell localization of candidate signatures

| **Signature** | **Comparison** | **Paired patients** | **Median difference** | **95% CI** | **Raw p** | **Holm p** |
| --- | --- | --- | --- | --- | --- | --- |
| Glutamine | Malignant vs T cell | 14 | 0.424 | 0.320–0.537 | 0.002 | 0.010 |
| Glutamine | Malignant vs NK cell | 9 | 0.398 | 0.151–0.673 | 0.009 | 0.037 |
| Glutamine | Malignant vs B cell | 11 | 0.454 | 0.257–0.561 | 0.004 | 0.019 |
| Glutamine | Malignant vs Macrophage | 11 | 0.215 | 0.117–0.268 | 0.045 | 0.136 |
| Glutamine | Malignant vs Endothelial | 7 | 0.214 | -0.069–0.409 | 0.108 | 0.217 |
| Glutamine | Malignant vs CAF | 9 | 0.021 | -0.086–0.277 | 0.477 | 0.477 |
| IFN-γ | Malignant vs T cell | 14 | -0.356 | -0.442–-0.228 | 0.001 | 0.007 |
| IFN-γ | Malignant vs NK cell | 9 | -0.217 | -0.502–-0.013 | 0.024 | 0.098 |
| IFN-γ | Malignant vs B cell | 11 | -0.073 | -0.230–0.029 | 0.083 | 0.249 |
| IFN-γ | Malignant vs Macrophage | 11 | -0.806 | -1.020–-0.617 | 0.005 | 0.025 |
| IFN-γ | Malignant vs Endothelial | 7 | -0.125 | -0.346–0.052 | 0.353 | 0.353 |
| IFN-γ | Malignant vs CAF | 9 | -0.142 | -0.481–0.116 | 0.124 | 0.249 |

*Note:* Positive differences indicate higher scores in malignant cells. Scores were summarized per patient and compartment before paired testing; cells were not treated as independent replicates.

**Table 5.** Exploratory pretreatment response models in GSE91061

| **Model** | **Term** | **OR/SD** | **95% CI** | **p** |
| --- | --- | --- | --- | --- |
| additive | Glutamine | 0.697 | 0.322–1.405 | 0.319 |
| additive | IFN-γ | 1.135 | 0.564–2.351 | 0.722 |
| exploratory interaction | Glutamine | 0.732 | 0.320–1.598 | 0.427 |
| exploratory interaction | IFN-γ | 1.017 | 0.436–2.210 | 0.967 |
| exploratory interaction | Interaction | 0.680 | 0.279–1.426 | 0.323 |

*Note:* Firth penalized logistic regression; n=49 with 10 responders. OR, odds ratio. AUC values were calculated from leave-one-out predictions.

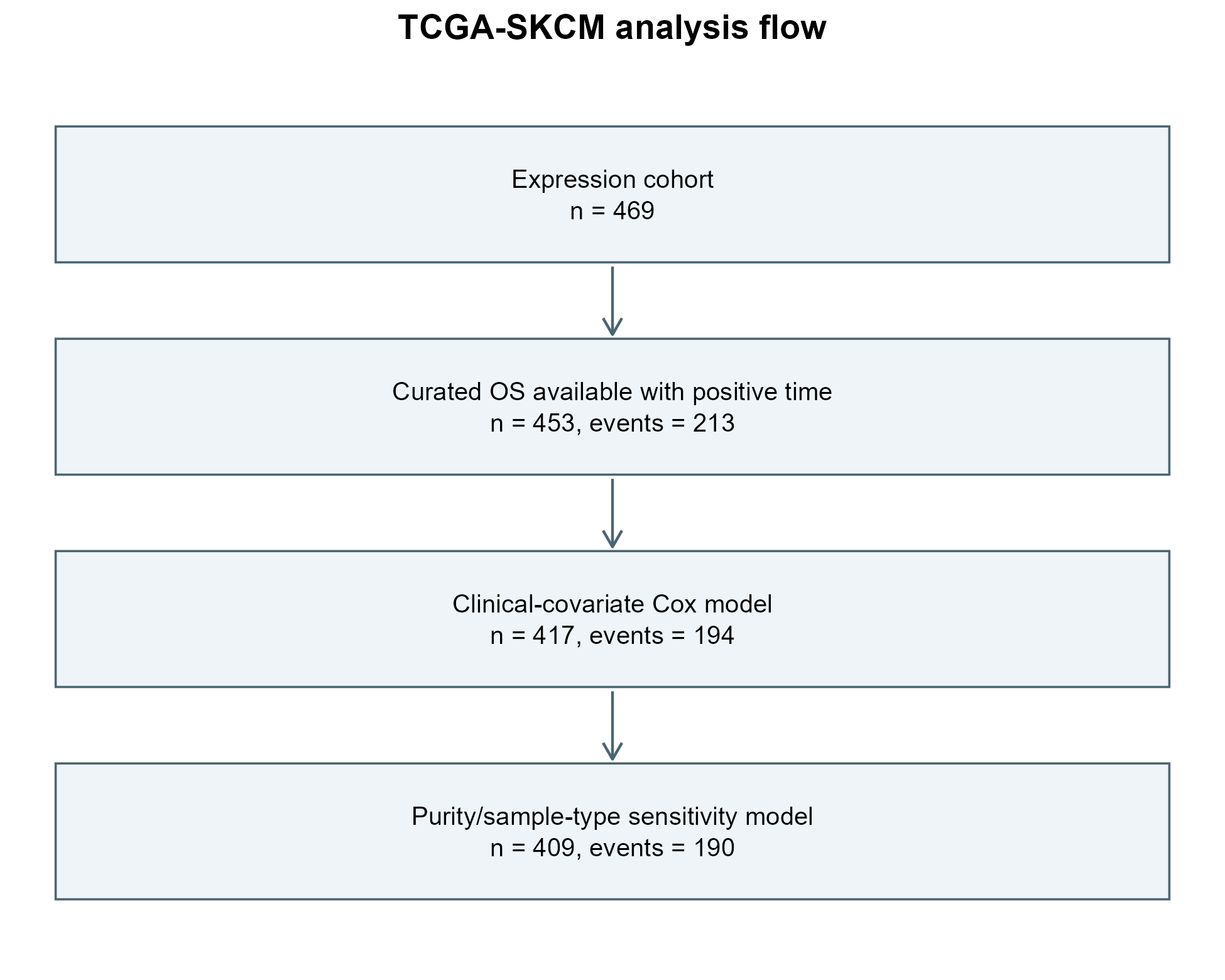

**Figure 1.** TCGA-SKCM analysis flow from aligned expression profiles to the overall-survival, complete-covariate, and composition-adjusted analysis sets. Table 1 reports the complementary external cohorts.

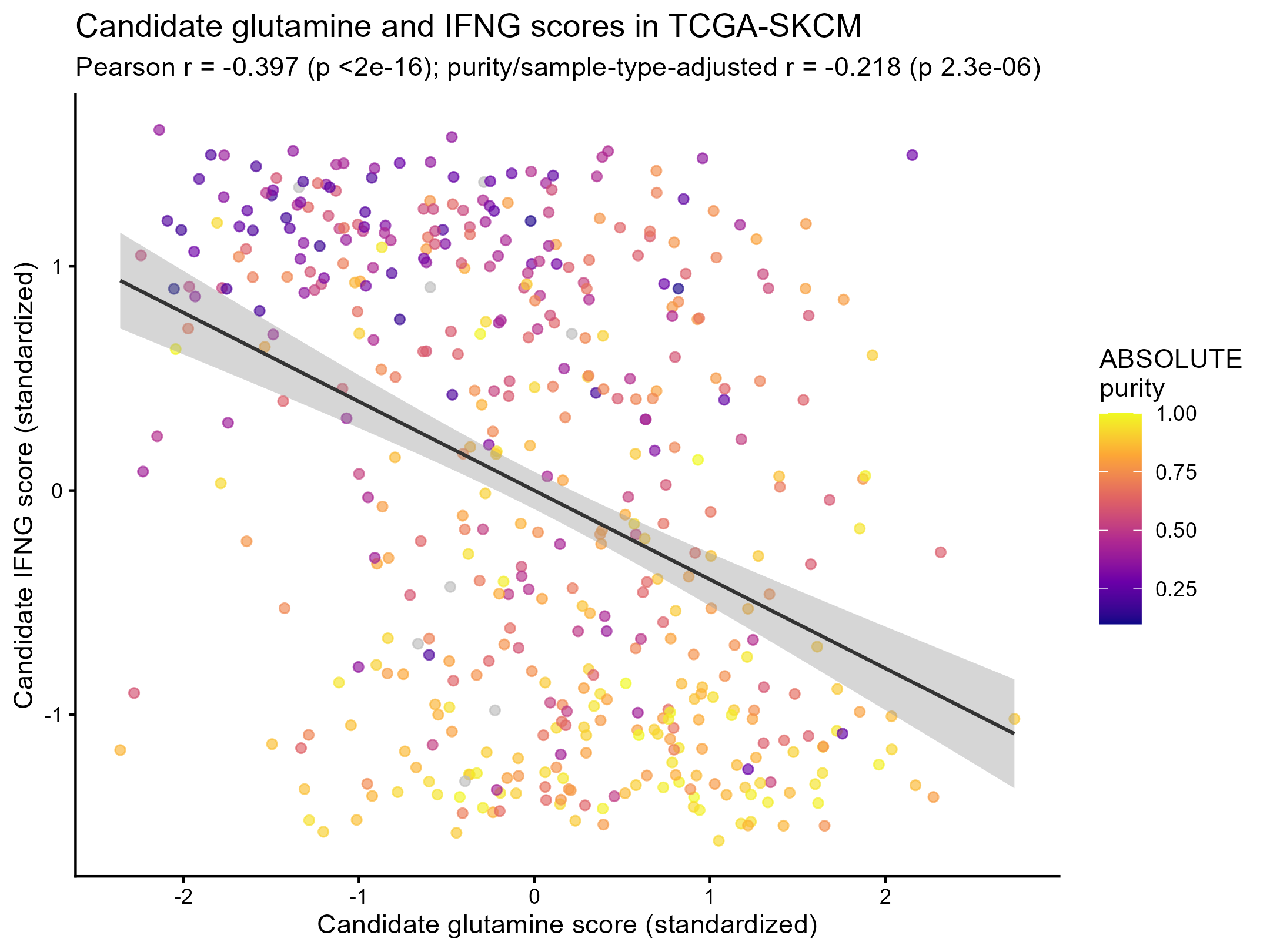

**Figure 2.** Inverse association between continuous candidate glutamine-associated and IFN-γ-associated GSVA scores in TCGA-SKCM. The line is the least-squares fit; the shaded band is its 95% confidence interval. This unadjusted bulk association is not interpreted as cell-intrinsic antagonism.

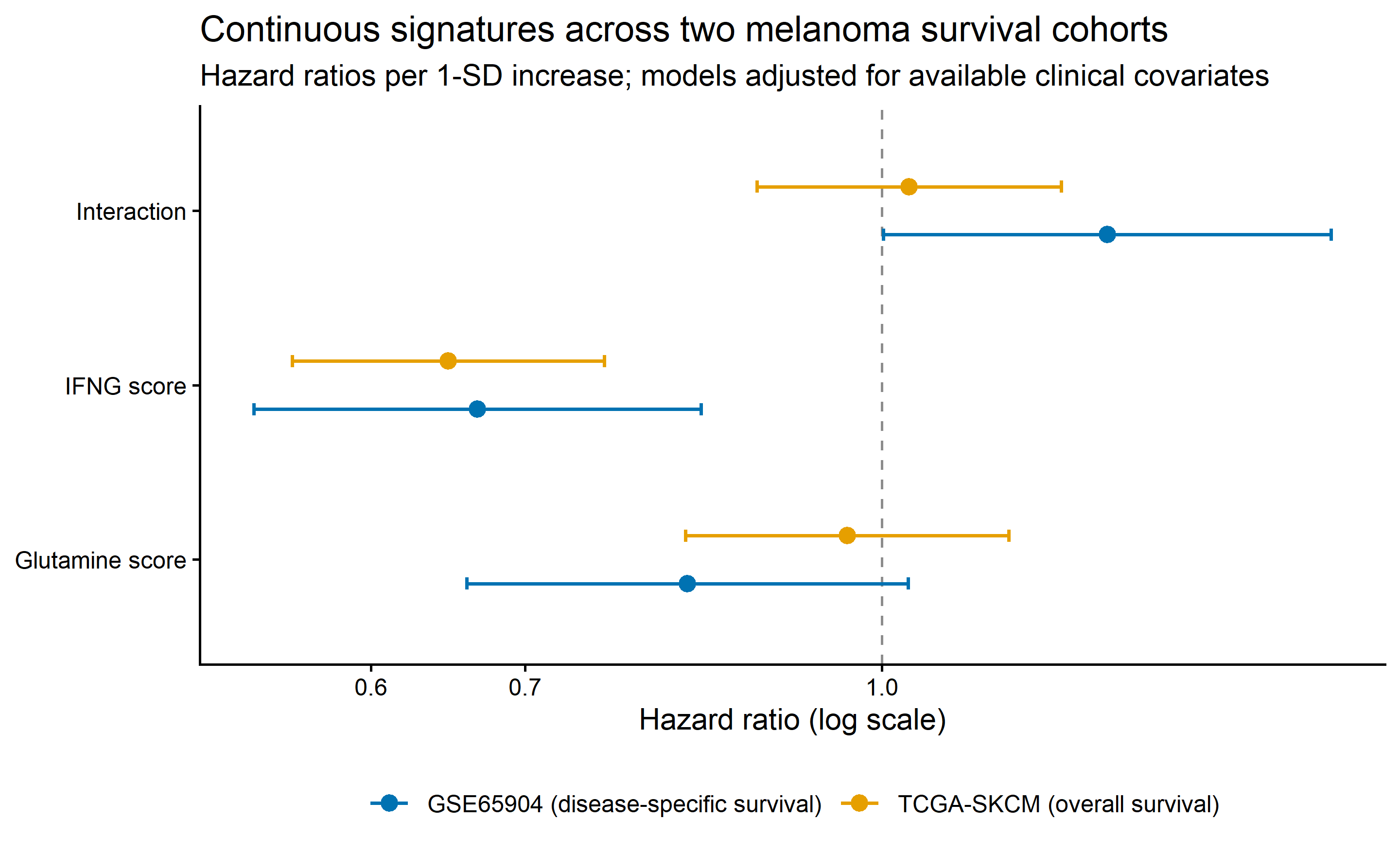

**Figure 3.** Cross-cohort adjusted survival associations per one-standard-deviation increase in score. IFN-γ-associated transcription is directionally consistent and statistically supported in TCGA OS and GSE65904 DSS. Glutamine and interaction estimates are not consistently supported. The dashed line marks HR=1.

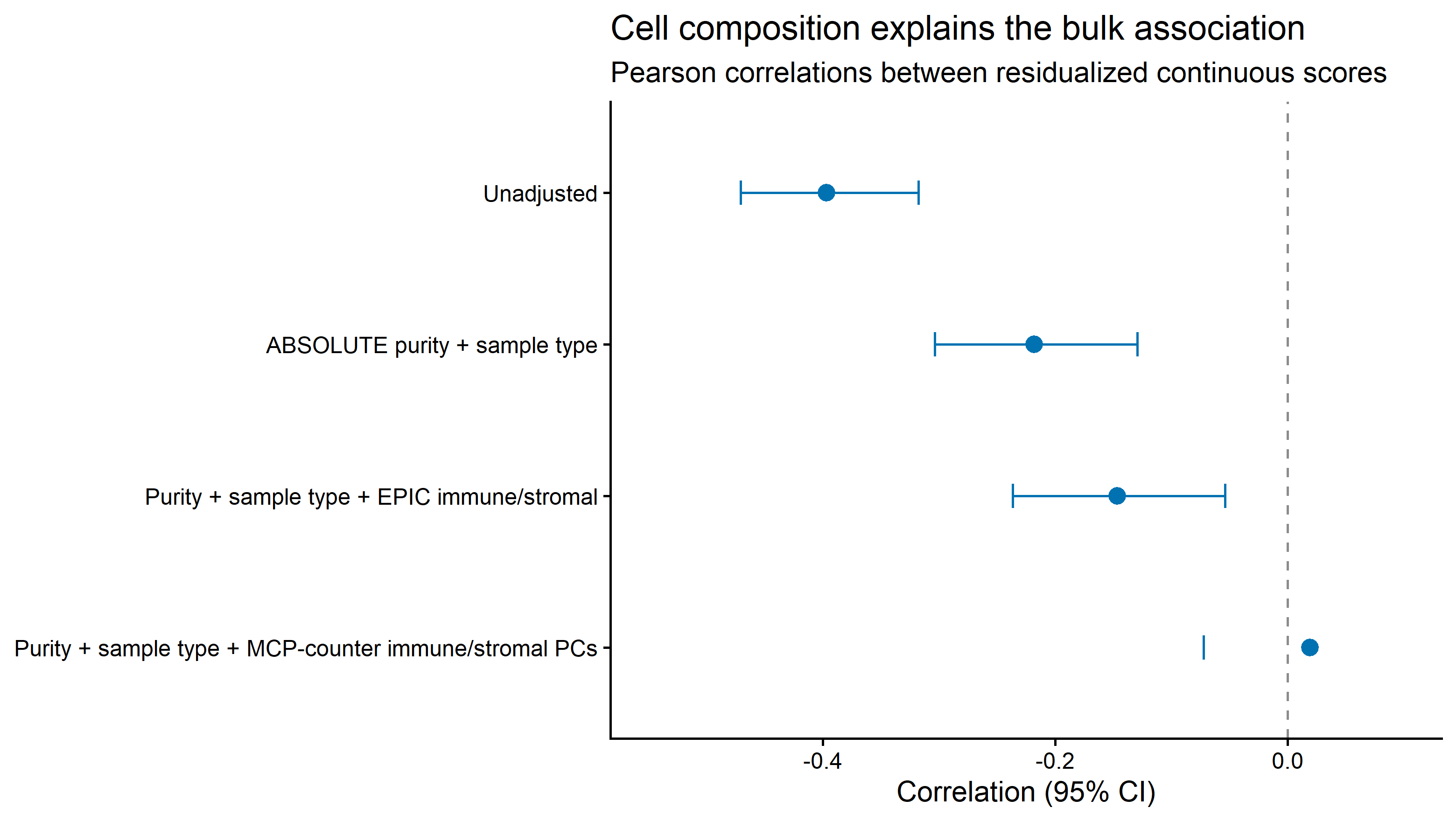

**Figure 4.** Progressive attenuation of the TCGA bulk glutamine–IFN-γ score correlation after adjustment for tumor purity, sample type, and immune/stromal composition. Points show Pearson correlations of residualized continuous scores; bars show 95% confidence intervals.

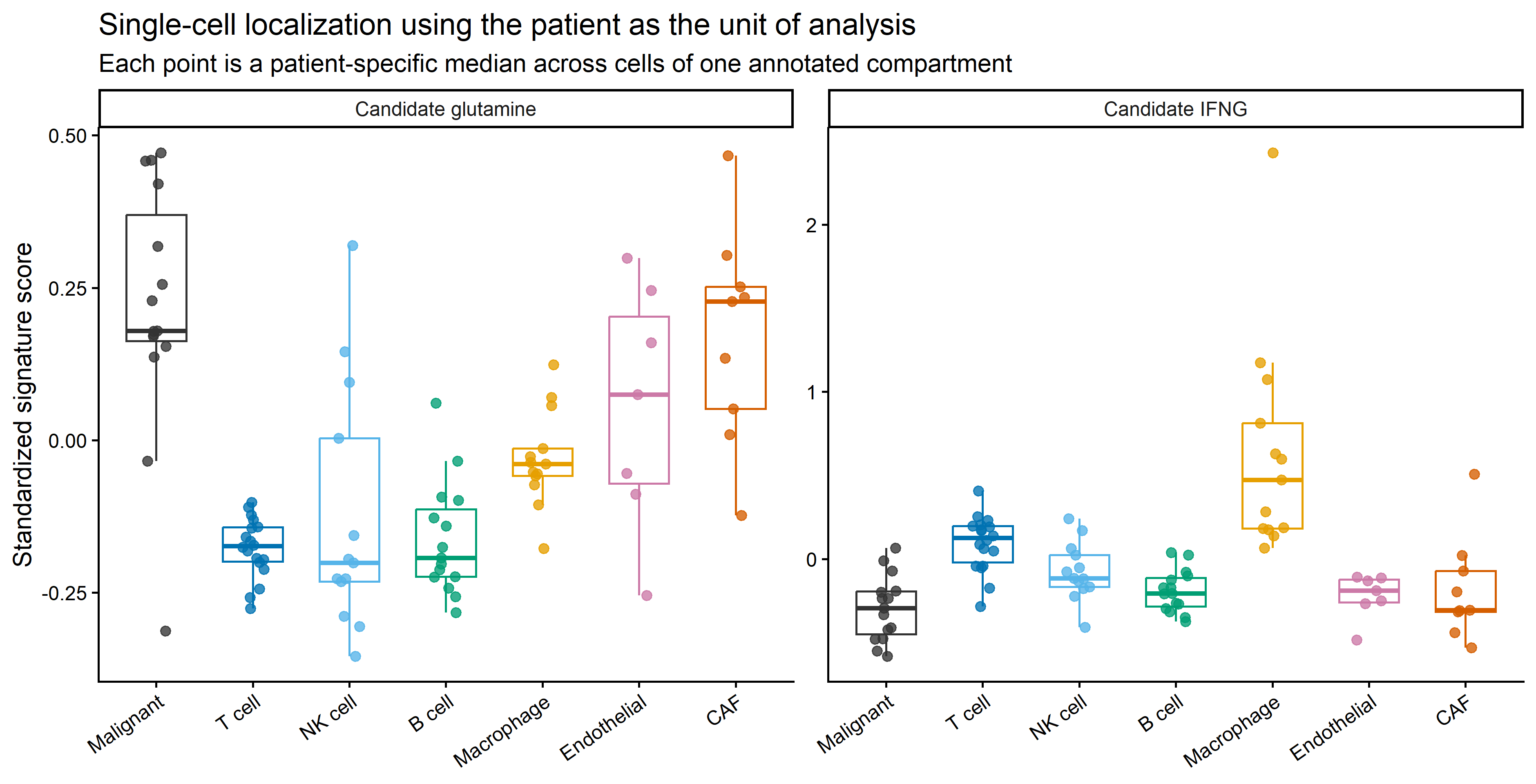

**Figure 5.** Patient-level single-cell localization in GSE72056. Each point is a patient-specific median score for one annotated compartment; boxplots summarize patients, not cells. Candidate glutamine-associated scores are relatively enriched in malignant cells, whereas IFN-γ-associated scores are enriched in immune compartments.

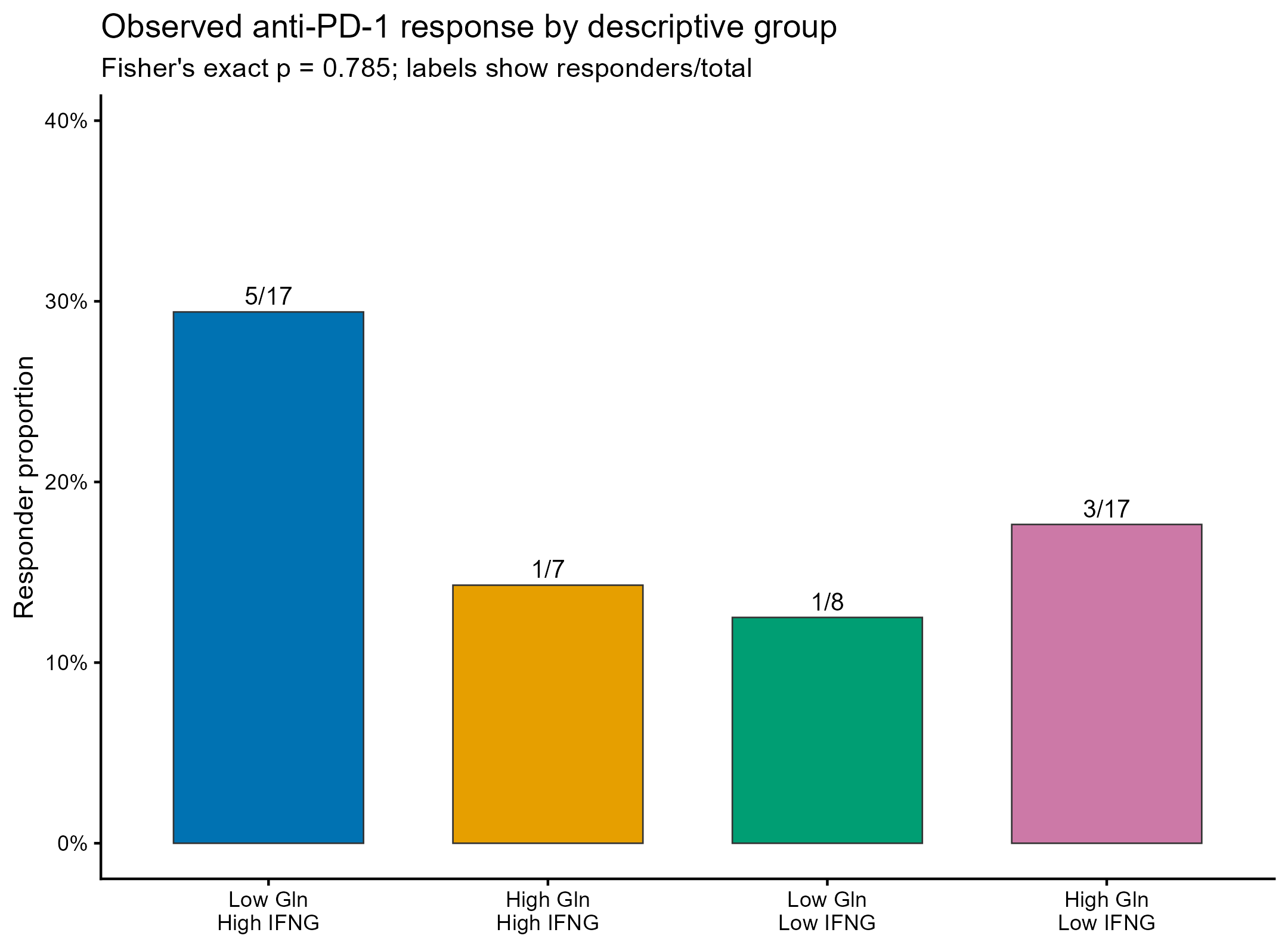

**Figure 6.** Exploratory pretreatment score distributions by response in GSE91061. The cohort contained 49 tumors and 10 responders. Firth regression and leave-one-out discrimination did not support a response association, but confidence intervals were wide.
